# Rodent-borne bacterial infections in Gwangju, Korea

**DOI:** 10.1101/411736

**Authors:** Mi Seon Bang, Choon-Mee Kim, Dong-Min Kim, Na Ra Yun

## Abstract

**Background:** This study investigated the prevalence of *Orientia tsutsugamushi, Anaplasma phagocytophilum,* and *Leptospira interrogan* in wild rodents through molecular detection in organ samples collected from two distinct sites in Gwangju Metropolitan City, Republic of Korea.

**Methodology/Principal Findings:** A total of 47 wild rodents, identified as *Apodemus agrarius (A. agrarius)*, were captured from June to August 2016. The seroprevalence of antibodies against bacterial pathogens in rodent sera was analyzed; 17.4% (8/46) were identified as *O. tsutsugamushi* through indirect immunofluorescence assay and 2.2% (1/46) were identified as *Leptospira* spp. through passive hemagglutination assay. Using molecular methods, the spleen, kidney, and blood samples were evaluated for the presence of *O. tsutsugamushi, A. phagocytophilum*, and *L. interrogans*. Out of 47 wild rodents, 25.5% (12/47) were positive for bacterial pathogens by PCR, where 19.1% (9/47) were positive for *A. phagocytophilum* and 6.4% (3/47) were positive for *L. interrogans*, while none were positive for *O. tsutsugamushi*. In addition, testing for bacterial infection in different tissues indicated that 8.7% (4/46) were positive for *A. phagocytophilum* in the blood, 13.3% (6/45) were positive for *A. phagocytophilum* in the spleen, and 6.4% (3/47) and 2.1% (1/47) were positive for *L*. *interrogans* and *A. phagocytophilum*, respectively, in the kidney.

**Conclusions/Significance:** In this study, tropisms for *A. phagocytophilum* in the spleen and *L. interrogans* in the kidney were identified. Notably, *A. phagocytophilum* and *L. interrogans* were detected in wild rodents living in close proximity to humans in the metropolitan suburban areas. Results of the present study indicate that rodent-borne bacteria may be present in wild rodents in the metropolitan suburban area of Republic of Korea.

**Author Summary:** Many zoonotic diseases are spreading not only in the Republic of Korea (ROK), but also worldwide. Scrub typhus, anaplasmosis, and leptospirosis are well known diseases that are considered common, widespread rodent-borne infectious diseases. Rodents serve as important reservoirs for zoonotic pathogens such as *O. tsutsugamushi, A. phagocytophilum*, and *L. interrogans,* which may be fatal to humans. Our study demonstrated the prevalence of these pathogens in wild rodents, through molecular assays and seroprevalence in organ samples. We captured 47 wild rodents in the Gwangju metropolitan city area of ROK. All were identified as *A. agrarius*. The prevalence of rodent-borne bacteria was 17.4% in the sera, where 25.5% was positively detected as bacterial pathogens via polymerase chain reaction. Our results indicate the importance of detecting rodent-borne bacteria in wild rodents living close to humans in suburban areas of ROK. Our data was limited to only a few samples of rodents in two regions. More samples may have to be collected over longer periods of time, to investigate the infectious nature of these pathogens in detail.

## Introduction

Rodents are known carriers of zoonotic pathogenic agents that are usually transmitted to humans through direct or indirect contact [1]. Among the known zoonotic diseases, scrub typhus, anaplasmosis, and leptospirosis are considered to be common, widespread rodent-borne infectious diseases.

Scrub typhus, which is caused by *Orientia tsutsugamushi*, is an infectious disease and has become one of the most prevalent human diseases in Republic of Korea (ROK). Since 2004, the disease has spread widely in the southwestern provinces of the country and is endemic [2]. In 1995, 274 cases of scrub typhus were reported, whereas 10,365 cases were reported in 2013, indicating a 38.1-fold increase in the incidence of this disease in ROK [3].

*Anaplasma phagocytophilum* causes human granulocytic anaplasmosis. *A. phagocytophilum* has been detected in *Hemaphysalis longicornis (H. longicornis), Ixodes nipponensis (I. nipponensis)*, and *Ixodes persulcatus (I. persulcatus)* ticks in ROK [4]. The incidence of anaplasmosis has also increased, from 1.4 cases per million in 2000 to 6.1 cases per million in 2010 [5, 6].

Leptospirosis has been one of the most important epidemic diseases in ROK since 1984. Wild rodents, particularly *A. agrarius*, are the common source of infection, especially during harvest season in rural areas of ROK. The prevalence of *Leptospira* infection in field rodents has been previously investigated via the detection of leptospiral DNA in rodent kidneys [7, 8].

The presence of scrub typhus, anaplasmosis, and leptospirosis in ROK indicates that *O.tsutsugamushi, A. phagocytophilum*, and *Leptospira interrogans* have infected both rodents and humans. Therefore, it is necessary to monitor the retention rate of pathogens in animal hosts, such as wild rodents, which harbor such vectors. A limited number of studies have been conducted on pathogen infection rates in wild rodents in the suburban areas of metropolitan cities. Especially, studies on the issue of organ tropism (i.e., the infection rates of *O. tsutsugamushi, A. phagocytophilum*, and *L. interrogans* in different tissues) are very limited.

This study investigated the prevalence of *O. tsutsugamushi, A. phagocytophilum*, and *L. interrogans* in wild rodents through molecular detection of these pathogens in organs of rodents collected from two distinct sites in Gwangju City, ROK. This study may broaden the understanding of health risks associated with wild rodents living in close proximity to humans.

## Methods

### Collection of wild rodent samples

The rodents were captured using Sherman traps (3” × 3.5” × 9’, USA) at two sylvatic habitats located in the suburban areas of Buk-gu and Gwangsan-gu in the Gwang-ju metropolitan city of ROK from June to August 2016. Captured rodents were numbered sequentially. All rodents were euthanized in accordance with an approved animal use protocol. Blood, spleen, and kidney tissues were obtained and stored at −80°C for use in future experiments. All samples were tested for the presence of *O. tsutsugamushi, A. phagocytophilum*, and *L. interrogans*.

### Detection of antibodies against bacterial pathogens in rodent serum

Blood samples were centrifuged at 3000 rpm for 20 min, following which serum was separated and maintained at 4°C. Indirect immunofluorescence assay (IFA) and passive hemagglutination assay (PHA) were performed 24–36 h after euthanasia.

For IFA, 10 µL of sera from each rodent was used in two-fold serial dilutions of 1:16 to 1:2048 in phosphate-buffered saline (PBS; pH 7.2; Welgene Inc., Korea). Diluted sera were deposited on antigen slides, incubated at 37°C for 30 min in a humidified chamber, washed twice with sterile PBS, and again washed twice with distilled water. A volume of 25 µL of fluorescein isothiocyanate (FITC)-conjugated goat anti-mouse IgG (Sigma, St. Louis, Missouri, USA) was added to each slide, incubated at 37°C for 30 min in a humidified chamber, washed twice with sterile PBS, washed twice with distilled water, and air-dried.

Mounting medium (Sigma) was added to each slide which was covered with a cover slip. The slides were examined for specific spots using a fluorescence microscope (Carl Zeiss, Oberkochen, Germany). A cutoff titer of 1:16 was used to determine seropositivity. Antigen slides for *O. tsutsugamushi* were provided by the Korea Center for Disease Control (KCDC). For PHA, Genedia Lepto PHA (Green Cross, Seoul, Korea) kit reagents were used. Serum (25 μL) from each rodent was added to a 96-well plate, and diluted to 1:80 in a dissolved solution of Genedia Lepto kit. Sheep blood (75 μL) containing red blood cells sensitized to *Leptospira* spp. was placed in the diluted serum for the agglutination assay. The Ag-Ab agglutination reaction at 1:80 dilution tested positive for leptospirosis.

### Molecular detection of bacterial pathogens in rodent tissues

A small piece of tissue (∼50 mg) in 200 μL of ATL buffer of the QIAamp Tissue & Blood Mini Kit (QIAGEN, Germany) was homogenized with the bead beater (BioSpec 3110BX Mini-Beadbeater-1 High-Energy Cell Disrupter). Tissue lysates were incubated at 56°C overnight, and genomic DNA was extracted using the QIAamp Tissue & Blood Mini Kit according to the manufacturer’s instructions.

To screen for *O. tsutsugamushi, A. phagocytophilum*, and *L. interrogans* DNA, nested PCR (nPCR) targeting each gene was performed using the listed primers (Table 1). nPCR was performed in 20 μL reaction volumes using the AccuPowerR PCR PreMix (Bioneer Corp., Korea). Each PCR mixture consisted of 16 μL of distilled water, 1 μL of each primer (10 pmol/μL), and 2 μL of DNA template. PCR was also performed using the INNOPLEX TSUTSU detection kit (cat. No. IPC10040; Intron Biotechnology). The kit contained primer sets designed to detect the 475-bp fragment of a gene encoding the 56 kDa antigen of *O.tsutsugamushi*. PCR was performed with 2 μL of DNA extract and 18 μL of distilled water treated with diethyl pyrocarbonate (DEPC; Gendepot, Barker, Texas, USA) in the PCR premix tube following the manufacturer’s instructions.

**Table 1.**
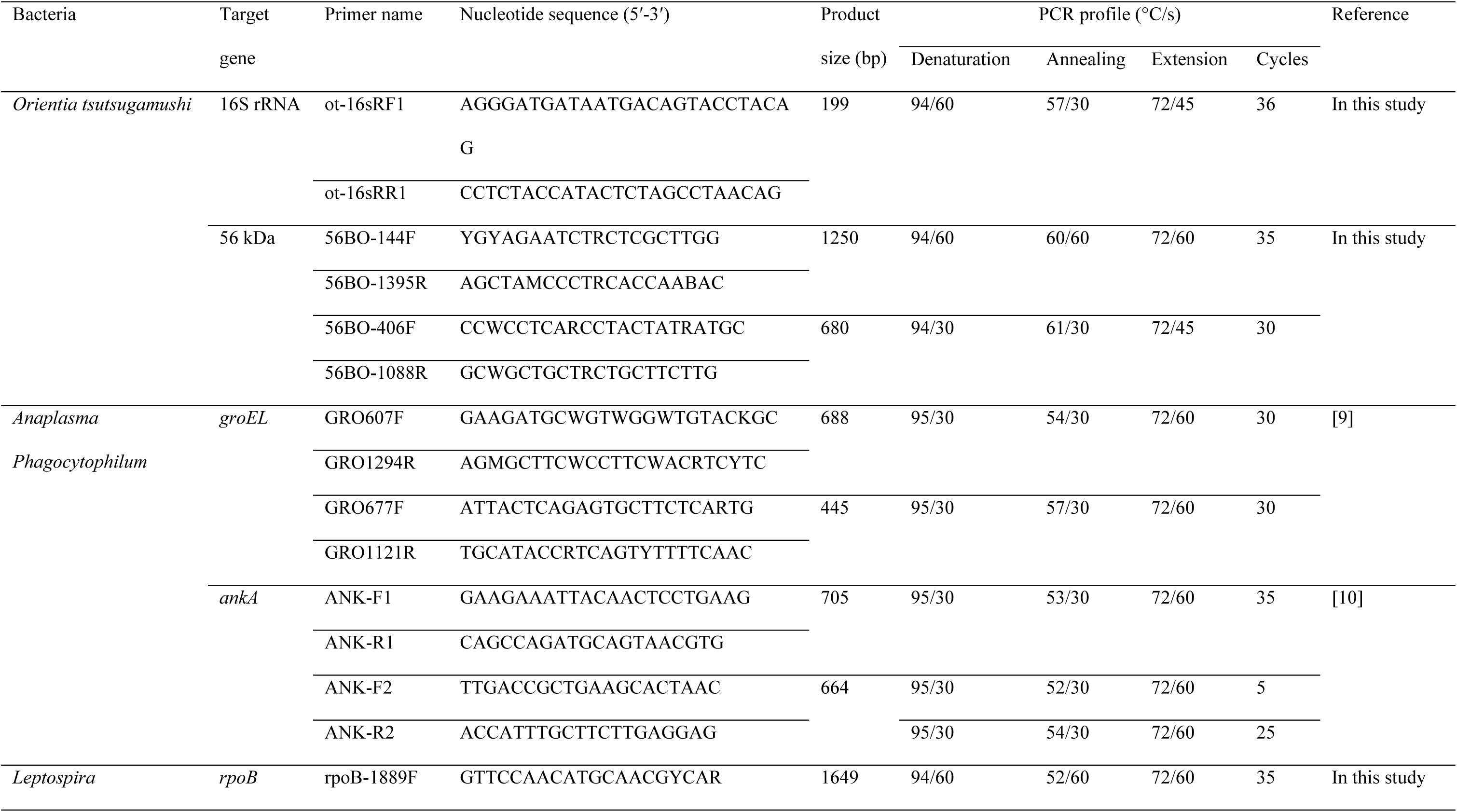
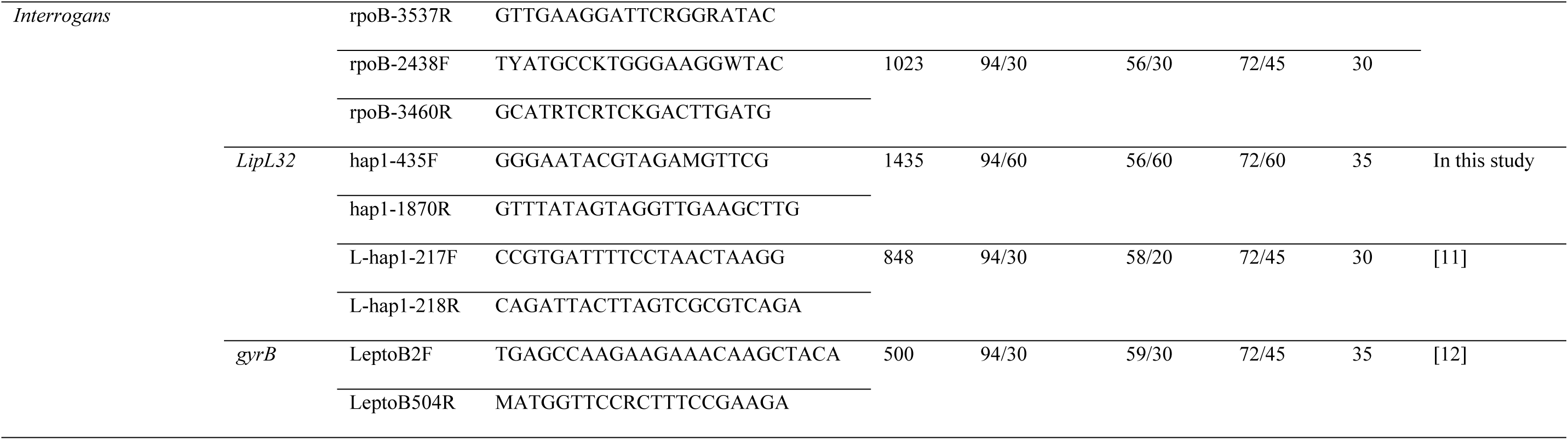
Nucleotide sequences and PCR conditions for the detection of rodent-borne bacteria in rodent tissues

PCR conditions and the amplicon sizes are listed (Table 1). All amplifications were carried out in an AB thermal cycler (Applied Biosystem, Foster City, CA, USA). Amplified products were separated by electrophoresis on a 1.2% agarose gel and stained with ethidium bromide for visualization.

### Ethics Statement

This study was approved from institutional review board (IRB) of Chosun University. All rodents were euthanized in accordance with an approved animal use protocol from Chosun University Institutional Animal Care and Use Committee (CIACUC) under approval number CIACUC2016-A0003.

### Sequencing and phylogenetic analysis

Sequencing of PCR amplicons from rodent-borne pathogens was conducted by Solgent Inc. (Daejeon, Korea). Obtained sequences were compared for similarity with sequences deposited in GenBank using BLAST. Gene sequences, excluding the primer regions, were aligned using the multisequence alignment program in Lasergene version 8 (DNASTAR, USA), and phylogenetic analysis was performed using the MEGA 6 software.

Phylogenetic trees were constructed using ClustalW of the MegAlign Program (DNASTAR, USA) based on the alignments of positive gene sequences using the neighbor-joining method. Bootstrap analysis (1,000 replicates) was carried out according to the Kimura 2-parameter method. Pairwise alignments were performed with an open-gap penalty of 10 and a gap extension penalty of 0.5.

## Results

### Wild rodent collection

A total of 47 wild rodents were captured from June to August 2016 at two trapping sites in the vicinity of Gwangju city in ROK. The wild rodents collected were: 16 mice captured in June; 18 mice captured in July; 13 mice captured in August. All wild rodents were identified to be *A. agrarius, w*hich were captured in fallow ground, around water, a boundary area between forest and field, and around a tomb.

### Seropositivity for *O. tsutsugamushi* and *L. interrogans* in rodent serum

Serum samples collected from 46 *A. agrarius* were subjected to IFA. Eight samples out of 46, (17.4%) were seropositive for *O. tsutsugamushi* with a cutoff titer ≥ 1:16 for IgG. In addition, PHA was performed for the detection of antibodies against *Leptospira* spp. and one sample out of 46 (2.2%) was positive with cutoff titer at 1:160 (Table 2).

**Table 2.** Rate of positive bacterial infections in 46 *A. agrarius* rodents, as indicated by serological assays

| Name of positive sera | Seropositive |  |
| --- | --- | --- |
|  | IFA titer for *O. tsutsugamushi* <sup>a</sup><br>(% positive rate) | PHA for *Leptospira* spp. <sup>b</sup><br>(% positive rate) |
| 6-6 | 1:128 | — <sup>c</sup> |
| 6-7 | 1:64 | — |
| 7-4 | 1:16 | — |
| 7-13 | 1:16 | — |
| 7-17 | 1:16 | 1:160 |
| 7-18 | 1:32 | — |
| 8-7 | 1:256 | – |
| 8-9 | 1:64 | – |
| **No. of positive sera** | **8 (17.4)** | **1 (2.2)** |

<sup>a</sup> cutoff titer of Immunofluorescence Assay, Immunoglobulin G $\geq 1:16$
<sup>b</sup> cutoff for Passive Hemagglutination Assay $\geq 1:80$
<sup>c</sup> –: negative

### Molecular detection of bacterial pathogens in rodent tissue

Tissue samples were obtained from the blood, spleen, and kidneys of 47 *A. agrarius*. Twelve out of 47 (25.5%) were positive for bacterial pathogens through PCR. Nine out of 47 (19.1%) were positive for *A. phagocytophilum*, while 3 out of 47(6.4%) were positive for *L.interrogans*. On the contrary, all rodent tissues were negative for *O. tsutsugamushi*.

Four blood samples out of 46 (8.7%) were positive for *A. phagocytophilum*. In spleen tissues, 6 samples out of 45 (13.3%) were positive for *A. phagocytophilum*, whereas none were positive for either *O. tsutsugamushi* or *L. interrogans*. In kidney tissues, 3 samples out of 47 (6.4%) were positive for *L. interrogans* and 1 sample out of 47 (2.1%) was positive for *A. phagocytophilum* (Table 3).

**Table 3.**
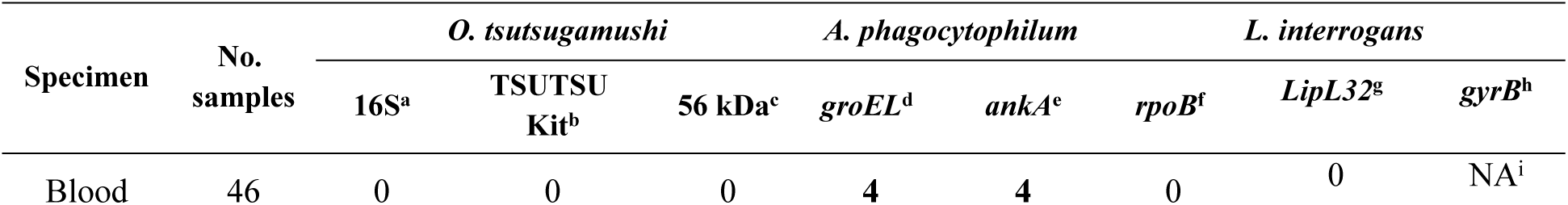
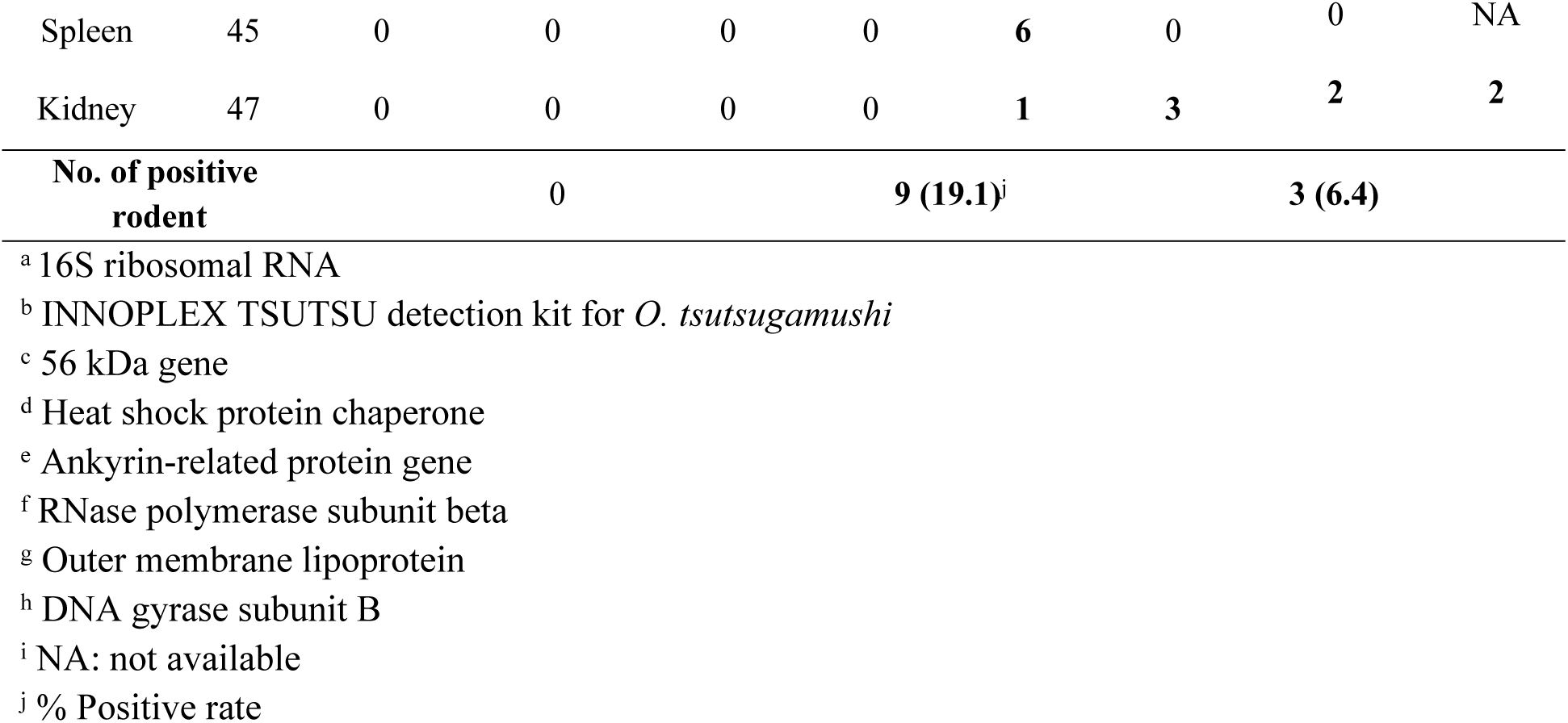
Number of positive detections for *Orientia tsutsugamushi, Anaplasma phogocytophilum*, and *Leptospira interrogans*, among the 47 *Apodemus agrarius* rodents, obtained via PCR targeting different genes

### Sequencing and phylogenetic analysis

Genes of rodent-borne bacteria were sequenced and aligned with sequences from the GenBank database using ClustalW. The neighbor-joining tree was constructed using the Kimura 2-parameter model (1,000 bootstrap replicates) in the MEGA 6 software.

The *ankA* gene sequences collected from rodent tissues demonstrated 99% similarity with that of *A. phagocytophilum* strain isolated from human in ROK (accession no. KJ677106 and KT986059, 100% bootstrap support, Fig 1A). The *groEL* gene sequences demonstrated 99% similarity with *A. phagocytophilum* strain isolated from rodents and dogs in ROK (accession no. KT192430 and KU519286, 93% bootstrap support, Fig 1B), and with *A. phagocytophilum* strain isolated from humans in Austria (accession no. KT454993, 96% bootstrap support, Figure 1B).

**Fig 1.**
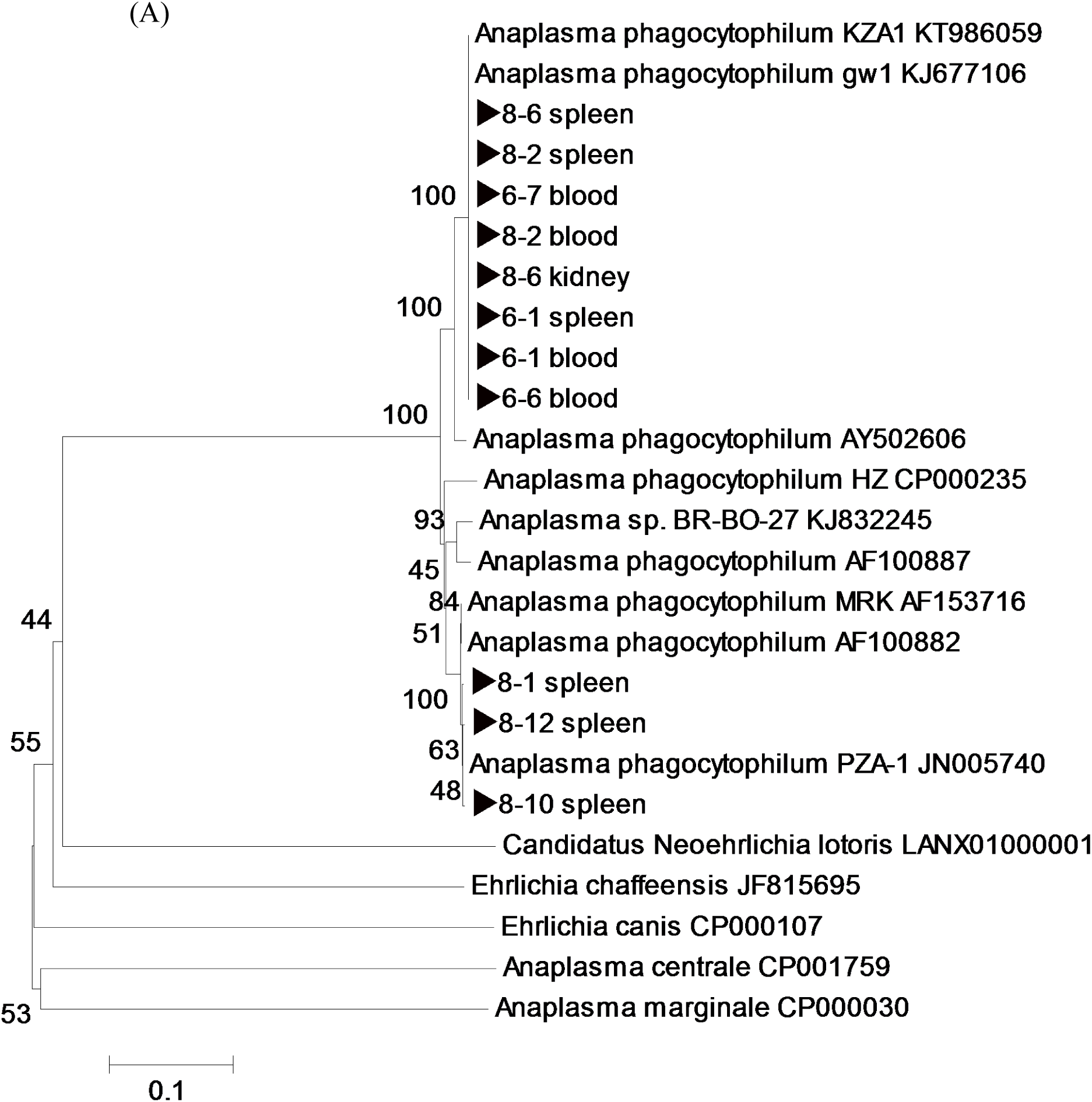
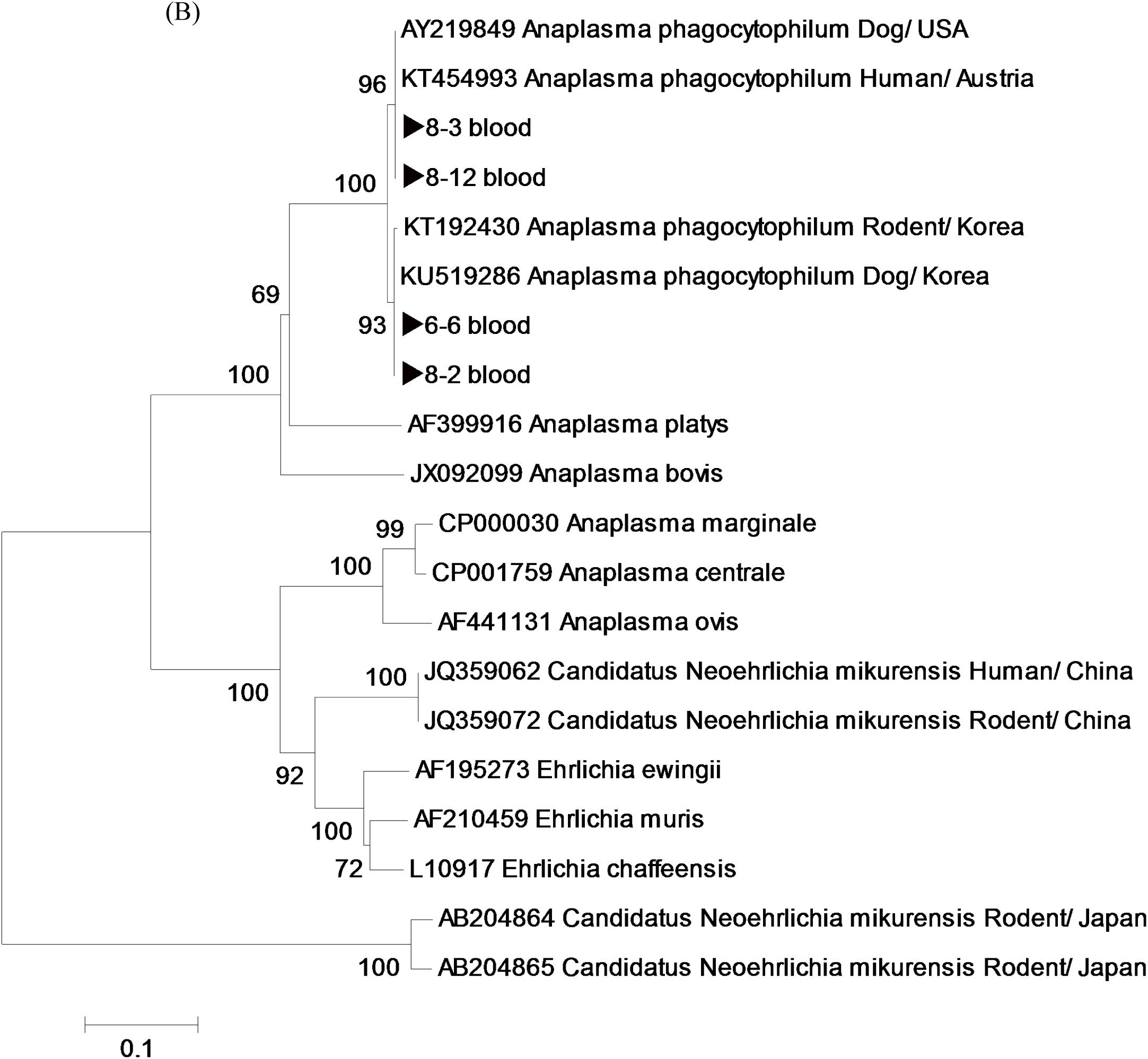
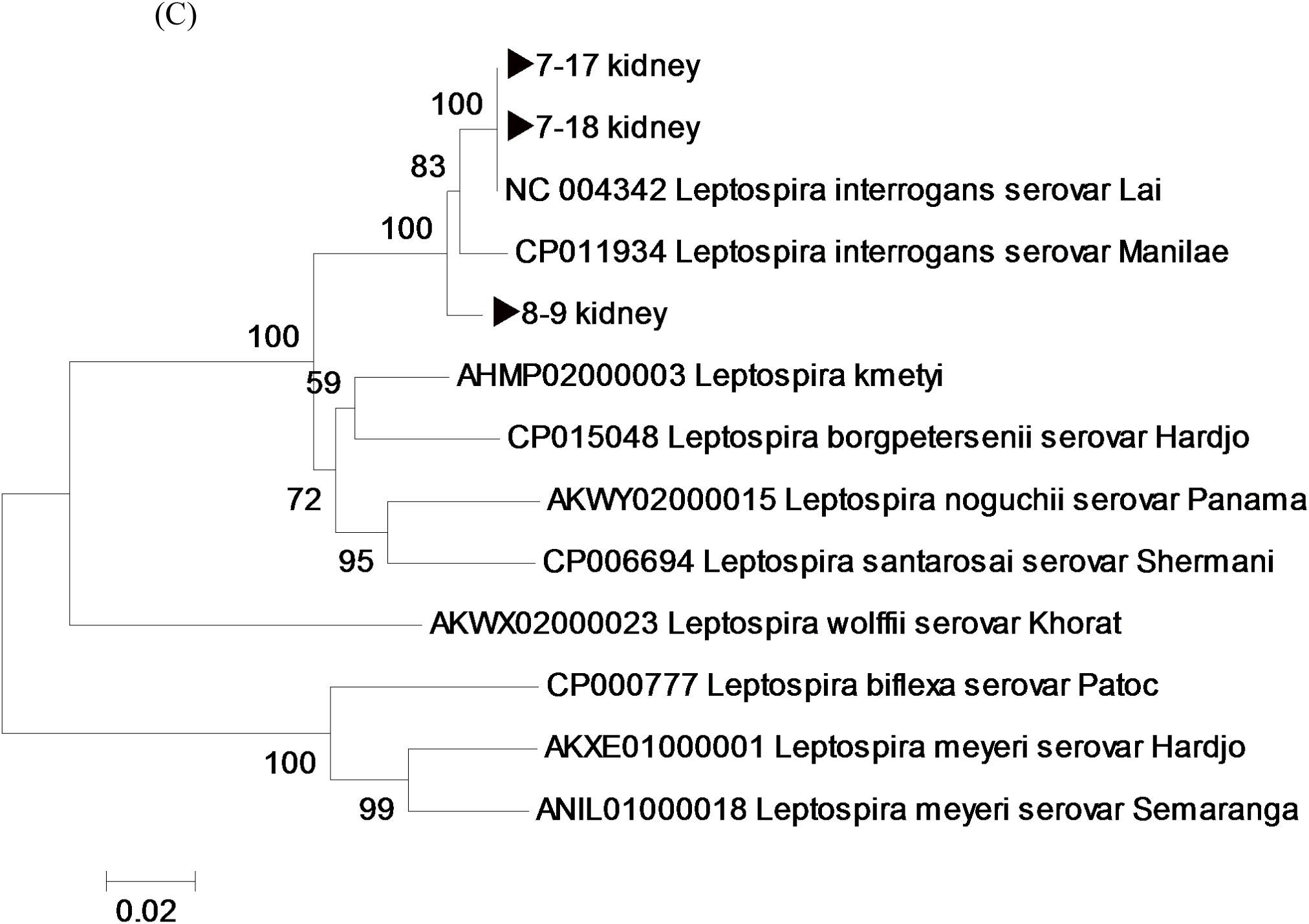
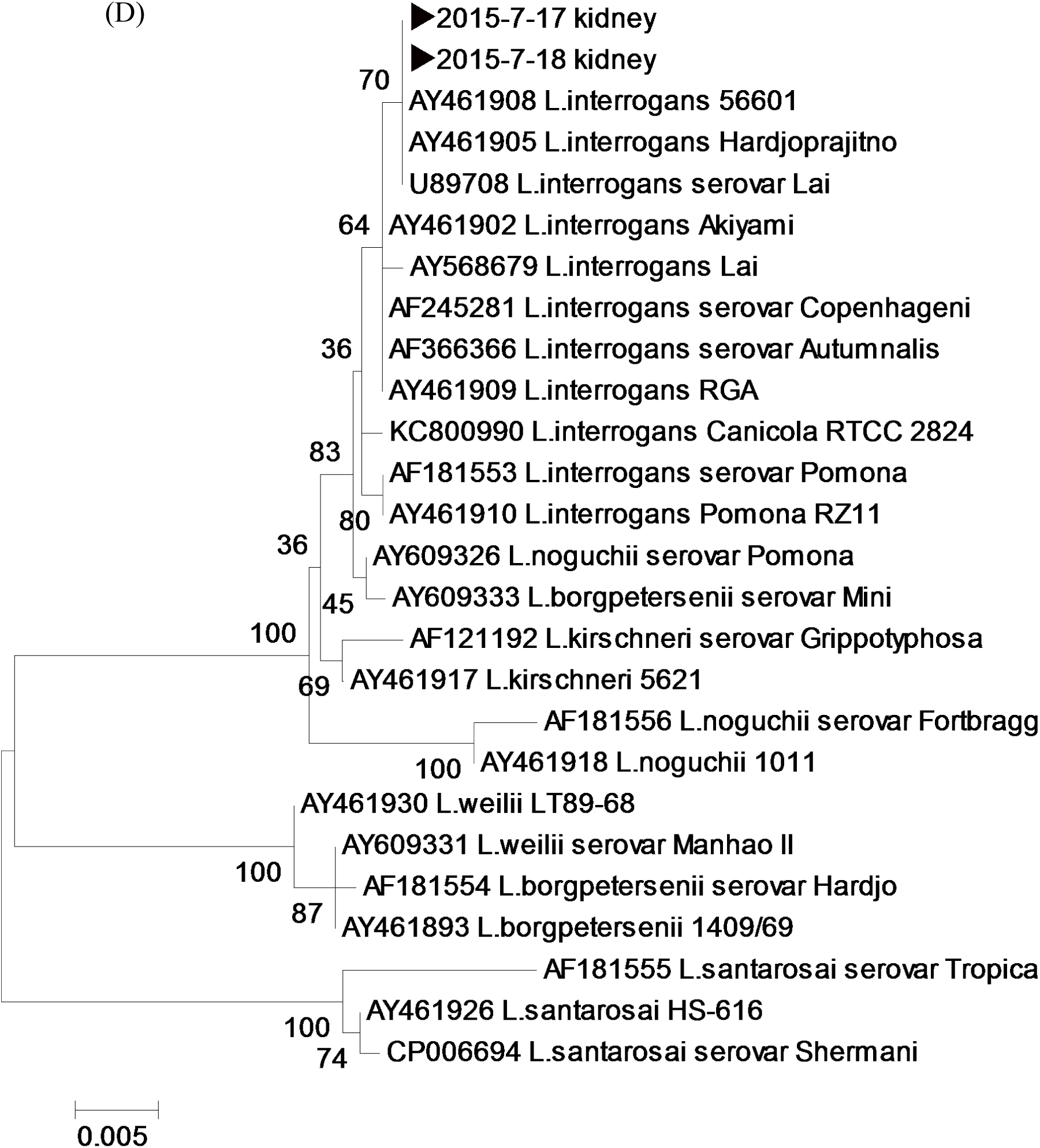
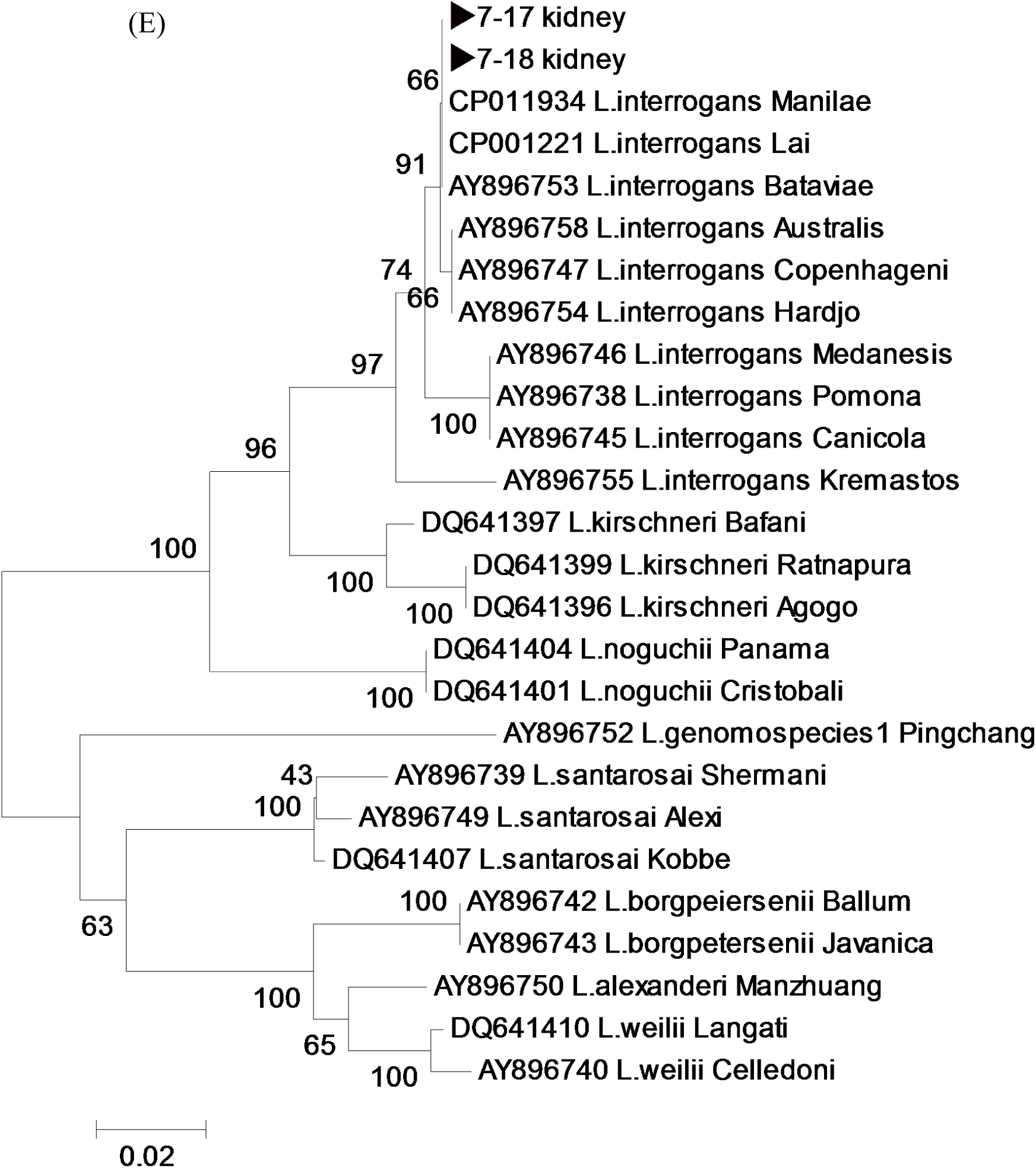
Phylogenetic trees generated based on nucleotide sequences of *A. phagocytophilum* and *L. interrogans* in tissues obtained from rodents captured in Gwangju. (A) 560 bp of *ankA* and (B) 330 bp of *groEL* gene sequences for *A. phagocytophilum* (C) 890 bp of *rpoB* gene, (D) 780 bp of *LipL32* gene, and (E) 400 bp of *gyrB* gene sequences for *L. interrogans*

The *rpoB* gene sequences collected from rodent tissues showed 99–100% similarity with *L. interrogans* serovar *lai* isolated from China (accession no. NC004342, 100% bootstrap support), (Fig 1C), while other samples exhibited 99–100% similarity with *L. interrogans* serovar *manilae* isolated from the mouse kidneys in Japan (accession no. CP011934, 100% bootstrap support), (Fig 1C). The *LipL32* gene sequences demonstrated 100% similarity with *L. interrogans* serovar *lai* isolated from sewage in China (accession no. AY461908) and with *L. interrogans* serovar *hardjoprajitno* isolated from humans in Indonesia (accession no.AY461905, 70% bootstrap support) (Fig 1D). The *gyrB* gene sequences demonstrated 100% similarity with *L. interrogans* serovar *lai* isolated from sewage in China (accession no. CP001221) and with *L. interrogans* serovar *manilae* isolated from mouse kidneys in Japan (accession no. CP011934, 66% bootstrap support), (Fig 1E).

### Discussion

Scrub typhus, which is caused by *O. tsutsugamushi*, is transmitted to humans via the bite of chigger mites (*Trombiculidae*) [13]. The incidence of scrub typhus is influenced by multiple factors, such as the behavior and population density of *Trombiculidae* and wild rodents. Human activity may also influence the infection rates [14, 15]. *A. agrarius* is a dominant wild rodent, which harbors chigger mites carrying *O. tsutsugamushi* [16]. Although, in this study, *O. tsutsugamushi* was not detected in any of the 47 *A. agrarius* tissues analyzed through PCR assays, they were positively detected in 8 of the 46 blood samples (17.4%) through IFA. The gold standard of assays for the serological detection of scrub typhus antibodies is the IFA [17]. Cutoff value for IgG in rodent sera in this study was 1:16. Kim et al. [18], observed that the IgG antibody titer increases abruptly over the first 2 weeks, reaching its peak at about 4 weeks in scrub typhus patients. Molecular techniques, such as PCR, provide the highest sensitivity and specificity for detection, especially in the early period of infection, due to the specificity of primers and low detection limits [19-23]. Our IFA and PCR results did not match, because it was unclear when these wild rodents had been infected. Thus, it may be preferable to perform both molecular and serological assays for proper pathogen detection.

*A. phagocytophilum* has long been recognized as an animal pathogen. However, it is now emerging as a human pathogen, which is of concern to public health [24]. This pathogen has been isolated from humans, as well as domestic and wild animals. The ticks carrying them have been found in the United States and in Europe [24-26]. In previous studies, serological and molecular evidence indicate that *A. phagocytophilum* has spread among humans, rodents, and ticks in many Asian countries, including Korea, China, and Japan [27-30].

Chae et al., [4], confirmed that ticks, rodents, and shrews near the Demilitarized Zone (DMZ) in Korea were positive for *A. phagocytophilum,* where 20 out of 403 rodent spleens (∼5%), tested positive for *A. phagocytophilum* in species-specific PCR assays. In the present study, 9 out of 47 *A. agrarius* (19.1%), were positive for *A. phagocytophilum*. Moreover, 6 out of 45 spleen samples, 4 out of 46 blood samples, and 1 out of 47 kidney samples were positive for *A. phagocytophilum.* These results displayed a 99% similarity with the results of those isolated from humans and dogs in ROK (Figs 1A and 1B). This suggests the possibility that wild rodents may be reservoirs of *A. phagocytophilum* in ROK.

*L. interrogans* is a highly motile, obligate aerobic spirochaete that causes leptospirosis in both humans and animals, including wildlife, livestock, and pets [31]. Rodents asymptomatically carry the bacteria in their kidneys and excrete them in urine, thus contaminating the environment [32, 33]. *L. interrogans* was detected in only 6.4% of the kidney samples in this study. In addition, PHA results indicate that only 1 out of 46 (2.2%) blood samples and only 1 of the 3 PCR-positive kidney samples were serologically positive for *Leptospira* spp. This suggests that the kidneys of wild rodents may be the main reservoirs of *L. interrogans*.

From the results of the phylogenetic tree analysis, the positive samples demonstrated 99– 100% similarity with serovar *lai* isolated from sewage in China and serovar *manilae* isolated from mouse kidney in Japan (Figs 1C-E). Since 1984, *L. interrogans* has been isolated from various sources, including human patients and wild rodents [34]. The *L*. *interrogans* isolates identified from *A. agrarius* field rodents were classified under serogroup Icterohaemorrhagiea and as serovar *lai* and *hongchon* [34].

Our results indicated the rate of rodent-borne bacterial infections in different tissues of 47 *A. agrarius,* collected from June to August 2016 in Gwangju City of ROK. A total of 12 (25.5%) wild rodents were positive for bacterial infection with *A. phagocytophilum* and *L. interrogans* as determined through PCR, and 8 (17.4%) sera samples were seropositive as determined through IFA and PHA. *A. phagocytophilum* and *L. interrogans* were detected in wild rodents living in close proximity to humans in suburban areas such as Gwangju City. In addition, higher levels of *A. phagocytophilum* and *L. interrogans* were detected in spleen and kidney tissues, respectively, compared to other tissues. In contrast, the positive rates detected in blood, using primer-specific PCR, were slightly lower. This signifies that overall rates of PCR detection can be lower if only blood samples were tested.

Although the seropositivity of *A. phagocytophilum* was not measured in this study, the detection rate of *A. phagocytophilum* in wild rodents using PCR was high.

The data were limited as only a few samples were obtained from only two sites. Future studies, using larger samples collected over longer periods of time may lead to a better understanding of the nature and spread of these infectious pathogens. It is felt that further research is required to monitor bacterial infection rates in wild rodents which serve as reservoir hosts for many infectious pathogens.

In conclusion, the present study indicates that rodent-borne bacterial infections circulating in wild rodent populations may be prevalent in the metropolitan suburban areas of ROK. In particular, tropisms of *A. phagocytophilum* for the spleen and of *L. interrogans* for the kidney were identified.

## Acknowledgments

Nothing

## Supporting information legends

Table S1. Number of positive detections among the 47 *Apodemus agrarius* assayed for bacterial infection through serological tests and PCR

